# A Common Agrochemical Mixture Increases Snail Hosts and Ecological Drivers of Schistosomiasis Risk

**DOI:** 10.64898/2026.01.12.698868

**Authors:** Marliece R. Barrios, Emily Selland, Meghan Forstchen, Jason Rohr

## Abstract

1. Agricultural intensification is accelerating across tropical regions, increasing fertilizer and pesticide inputs into freshwater ecosystems that regulate the transmission of many environmentally mediated diseases. For schistosomiasis, a major neglected tropical disease, transmission risk is strongly governed by the abundance of freshwater snail intermediate hosts and their aquatic habitat, yet most agrochemical studies have focused on single compounds rather than realistic mixtures.
2. Using replicated semi-natural aquatic microcosms, we experimentally tested how nitrogen– urea fertilizer and the widely used herbicide atrazine, applied alone and in combination at environmentally relevant concentrations, alter snail hosts (*Biomphalaria glabrata*), submerged aquatic vegetation (*Ceratophyllum demersum*), and algal resources that underpin schistosomiasis transmission.
3. Fertilizer and atrazine, both singly and together, increased snail habitat (*C. demersum*) and adult snail abundance. By reducing phytoplankton, atrazine likely alleviated shading on vegetation, thereby promoting habitat that supported snails through indirect bottom-up pathways. Agrochemical mixtures produced non-additive effects on host and habitat dynamics, indicating that transmission-relevant ecological responses cannot be predicted from single-compound exposures alone. Although parasite infection levels were low, the observed increases in snail hosts and habitat represent reliable ecological drivers of schistosomiasis risk.
4. *Synthesis and applications.* Our findings demonstrate that common agrochemical mixtures can restructure freshwater food webs in ways likely to elevate disease risk, highlighting the need to incorporate mixture-driven ecological responses into agrochemical risk assessment and agricultural management strategies in disease-endemic regions.

## INTRODUCTION

Agricultural expansion is one of the most pervasive drivers of global environmental change, reshaping freshwater ecosystems through increased nutrient and pesticide inputs (Tilman *et al*. 2011; Alexandratos & Bruinsma 2012). These chemical subsidies and stressors can profoundly alter aquatic food webs, with cascading consequences for biodiversity, ecosystem functioning, and human well-being (Clements & Rohr 2009; Rohr, Salice & Nisbet 2017). An applied ecological challenge of growing importance is understanding how agricultural pollution influences the transmission of environmentally mediated infectious diseases, where ecological processes rather than direct host–pathogen interactions often determine disease risk (Rohr *et al*. 2019; Hoover *et al*. 2020).

Schistosomiasis, a parasitic disease affecting over 250 million people worldwide, exemplifies this challenge (Gryseels *et al*. 2006; Steinmann *et al*. 2006; King 2010; McManus *et al*. 2018). Transmission depends on freshwater snails that serve as intermediate hosts for *Schistosoma* parasites. Despite the availability of the drug praziquantel, control is challenging because treatment does not prevent rapid reinfection when people return to snail-infested waters, reinforcing cycles of disease and poverty (King 2010; Doruska, Barrett & Rohr 2024). Human exposure to the parasite is tightly linked to ecological conditions that regulate snail abundance and habitat availability (Wood *et al*. 2019; Haggerty *et al*. 2020; Bakhoum *et al*. 2021a; Bakhoum *et al*. 2021b). Across endemic regions, particularly in sub-Saharan Africa, agricultural runoff has been repeatedly associated with increased snail populations, submerged aquatic vegetation, and parasite production (Hilali *et al*. 1985; Halstead *et al*. 2018; Haggerty *et al*. 2022; Haggerty *et al*. 2023; Rohr *et al*. 2023). As a result, ecological drivers such as snail host density and habitat extent are now recognized as strong predictors of schistosomiasis risk, often exceeding the predictive power of infection prevalence alone, especially in early or low-intensity transmission settings (Civitello *et al*. 2018; Wood *et al*. 2019; Civitello *et al*. 2022).

Fertilizers and pesticides play central roles in shaping these ecological pathways. Nutrient enrichment stimulates primary production and the growth of submerged aquatic vegetation, such as *Ceratophyllum demersum*, which provides both food and refuge for snail hosts and is positively associated with schistosomiasis transmission risk in both experiments (Best 1981; Haggerty *et al*. 2020; Rohr *et al*. 2023) and across sub-Saharan Africa (Chu 1978; Hilali *et al*. 1985; Madsen, Coulibaly & Furu 1987; Erko, Tedla & Petros 1991; Ofoezie 1999; Van Bocxlaer *et al*. 2011; Yirenya-Tawiah *et al*. 2011; Haggerty *et al*. 2020). In parallel, certain herbicides can indirectly promote snail populations by suppressing phytoplankton, increasing water clarity, and enhancing periphyton and macrophyte growth through bottom-up trophic pathways (Rohr *et al*. 2008; Staley, Rohr & Harwood 2011; Rohr, Halstead & Raffel 2012; Staley, Harwood & Rohr 2015; Rumschlag *et al*. 2019; Rumschlag *et al*. 2020). Experimental and field studies have shown that individual agrochemicals can elevate snail abundance and parasite transmission potential through these mechanisms (Rohr *et al*. 2008; Schotthoefer *et al*. 2011; Halstead *et al*. 2018; Becker *et al*. 2020; Haggerty *et al*. 2022).

Despite this growing body of work, agrochemicals are rarely introduced into aquatic ecosystems in isolation. In agricultural landscapes, fertilizers and pesticides typically co-occur as complex mixtures, even when applied separately to fields (Altenburger *et al*. 2013; Halstead *et al*. 2014). Ecological theory and empirical evidence demonstrate that such mixtures can interact in non-additive ways, producing outcomes that cannot be predicted from single-compound studies alone (Clements & Rohr 2009; Relyea 2009; Halstead *et al*. 2014). Nevertheless, most agrochemical risk assessments continue to rely on single-chemical evaluations, potentially mischaracterizing the ecological consequences of agricultural intensification for disease transmission (Boone *et al*. 2014; Rohr, Salice & Nisbet 2016; Rohr, Salice & Nisbet 2017).

From an applied ecology perspective, identifying how common agrochemical mixtures restructure disease-relevant food webs is therefore critical for designing management strategies that balance agricultural production with ecosystem and public health goals (Rohr *et al*. 2019; Hoover *et al*. 2020). This is particularly urgent in schistosomiasis-endemic regions, where fertilizer and pesticide use are projected to increase two- to five-fold in coming decades (Tilman *et al*. 2011; Alexandratos & Bruinsma 2012), and where small ecological changes can translate into large shifts in human disease risk (Rohr *et al*. 2023).

Here, we use replicated semi-natural aquatic microcosms to experimentally test the individual and combined effects of nitrogen–urea fertilizer and the widely used herbicide atrazine on key ecological drivers of schistosomiasis transmission. We focus on nitrogen–urea fertilizer because it accounts for the majority of fertilizer use across Africa (Research 2017), and on atrazine because it is among the most commonly applied herbicides in schistosomiasis-endemic regions (Dabrowski, Shadung & Wepener 2014) and is known to commonly influence aquatic primary producers and disease risk (Rohr & McCoy 2010; Rohr 2021). Specifically, we examine how these agrochemicals, applied at environmentally relevant concentrations, affect snail intermediate host abundance (*Biomphalaria glabrata*), submerged aquatic vegetation (*Ceratophyllum demersu*m), and algal resources. We hypothesized that fertilizer would increase snail abundance by enhancing bottom-up resource availability, that atrazine would indirectly promote snail habitat by reducing phytoplankton shading, and that agrochemical mixtures would generate non-additive effects on host and habitat dynamics.

By experimentally isolating these mechanisms in semi-natural systems, our study provides applied ecological insight into how realistic agrochemical mixtures restructure freshwater ecosystems in ways that are likely to influence schistosomiasis risk. More broadly, our results highlight the need to incorporate mixture effects into ecological risk assessment and agricultural management in disease-endemic landscapes.

## METHODS

### Experimental Design and Community Composition

We established 20 semi-natural aquatic microcosms in 20-L plastic aquaria in a laboratory at the University of Notre Dame (41°42′34.9′′N 86°14′16.0′′W) to examine the effects of agrochemical pollution (applied singly or in combination) on snail host populations, snail infection status, snail food resources, and vegetation biomass. Three weeks before agrochemical application, each tank was established in accordance with the Walstad Tank Method for formation of self-sufficient aquatic communities (Walstad 2013). Tanks were filled with 1-inch pre-washed play sand (Quikrete Play Sand) over 1-inch pre-sifted organic garden soil (Nature’s Care 1.5 cu. ft. Organic Garden Soil), followed by 12 L of artificial freshwater (COMBO). Water in tanks was allowed to age for 24 h before tanks were seeded with 1 L of water acquired from the on-campus lake (not filtered for periphyton, phytoplankton, or zooplankton) and 49.495±0.328 g *Ceratophyllum demersum* (sourced from Play It Koi, Botnell, Washington, USA), a submerged aquatic plant species that serves as habitat for *Schistosoma* harboring snails and is positively associated with human *Schistosoma* infections (Wood *et al*. 2019; Haggerty *et al*. 2020; Rohr *et al*. 2023). Tanks were placed directly underneath full spectrum lights set to a 12:12 hour light:dark cycle. All four walls of each tank were covered with black plastic so that light only entered from the tops of the tanks. Bubblers were added to the tanks to ensure that oxygen levels were sufficient to sustain snail populations. Over the three-week establishment period, water was mixed among tanks twice a week in an attempt to homogenize starting communities.

Before adding *C. demersum* to each tank, it was placed in a 10% bleach solution for 10 seconds before thoroughly rinsing it with reverse osmosis (RO) water and subsequently soaking the plants in 0.25 mg/L niclosamide solution for 24 hours. This eliminated any snails or macroinvertebrates on the *C. demersum*. The *C. demersum* was washed twice with RO water and then cut into 6-inch segments. Segments were spun dry with a salad spinner and dry weight was recorded before being added to the tank.

Immediately before agrochemical application (Week 0), each tank received six adult *Biomphalaria glabrata* [NMRI strain] snails (9.073±0.562 mm). These snails were obtained from a laboratory colony originally seeded with snails from the NIAID Schistosomiasis Resource Center (Biomedical Research Institute, Rockville MD, USA).

We randomly assigned one of four treatments to each of the 20 experimental tanks (*n* = 5): solvent control (0.0625 mL/L acetone), atrazine, nitrogen-urea fertilizer, and atrazine plus nitrogen-urea fertilizer (i.e., a 2 x 2 experimental design). Tanks with herbicide treatment were dosed twice, in Week 0 and Week 3, with an ecologically relevant dose of 100 ppb of atrazine (based on previous calculations using the US Environmental Protection Agency’s GENEEC v.2 software). The atrazine was dissolved in 500 mL acetone. Tanks with fertilizer treatment were dosed once in Week 0 with 3300 µg/L of nitrogen-urea fertilizer, a concentration similar to those found in waterbodies in the field (Chase 2003).

*Schistosoma mansoni* (NMRI strain) eggs were added directly to all tanks on Weeks 0, 2, and 4. *Schistosoma mansoni* eggs were extracted from livers of Swiss-Webster mice seven weeks after they were infected by the NIAID Schistosomiasis Resource Center. The eggs from two livers were maintained on 1.2% saline until they were evenly distributed among 25 5-mL aliquots. Each of the 20 tanks received one of these aliquots on each of the three dosing weeks and the five extra aliquots were used to estimate egg viability (i.e., how many miracidia hatched from the eggs in the aliquot). Egg viability was determined to be 1450.4, 940.8, and 1141.2 miracidiae hatched per 5 ml aliquot for Week 0, Week 2, and Week 4, respectively.

### Experimental Response Measurements

Adult snail populations were quantified weekly throughout the 19-week experiment beginning one week after the treatments were added. All snails were removed from the tank using a small net (10.4 x 7.5 x 8.5 cm) and counted if the adult population (adult snails were determined to be those >2mm in diameter) was <24. If the adult snail population in a tank visually exceeded 24 snails, snails were only collected from one randomly selected quarter of the tank and then scaled up to the entire tank (i.e., by multiplying by four). Snails on vegetation or tank walls within subsampled regions were picked off and included in counts if they exceeded 2mm in size.

Juvenile snail populations (snails identified to be <2mm in diameter) and snail egg masses were quantified weekly beginning in Week 7. One to two tank walls (26.67 x 30.48 cm) were selected randomly to count juvenile snails and egg masses. Population counts of adult and juvenile snails continued for the remaining duration of the experiment. At the end of the experiment (Week 19), we emptied the tanks and counted the snails in each tank and preserved live snails in 70% ethanol. We determined snail infection status under a dissecting microscope by inspecting the gonads of each preserved snail for *S. mansoni* after staining with Raillet-Henry’s solution at the end of the experiment (Théron, Pages & Rognon 1997).

In addition to quantifying snails, we also quantified photosynthetic taxa in each tank. Starting at Week 1, *C. demersum* was removed weekly from each tank, cleaned of snails, spun dry using a salad spinner, weighed, and returned to each tank. Periphyton and phytoplankton measurements were taken during Weeks 8, 12, 16, and 19 of the experiment. To measure periphyton, a single microslide (75 x 25 x 1mm) was suspended on each of the four walls of the tank at a depth of 15 cm. An ethanol-sterilized, COMBO-rinsed razor blade was used to scrape the periphyton from the bottom inch of the slide (pre-marked before microslide deployment) into a labeled beaker with 15 mL COMBO water and homogenized. A separate 960 µL water sample was collected from 5 cm below the surface of the water at the center of each tank for phytoplankton quantification. For each of these phytoplankton and periphyton samples, chlorophyll-a and photosynthetic efficiency were measured three times using a handheld AquaPen fluorometer (Z985 Cuvette AquaPen, Qubit Systems Inc., Kingston, Ontario, Canada) and then averaged.

Starting at Week 7, we measured cercarial production weekly for each sampled snail greater than 2mm in diameter (from the subsample removed from each tank for snail population size quantification). Each snail was placed under fluorescent light in an individual 30-mL beaker filled with 15 mL of COMBO. After 90 minutes, each snail was removed from their beaker using forceps and placed back into their respective tanks. Each beaker then received four drops of Lugol’s solution to stain and preserve any *S. mansoni* cercariae shed into the beaker. Cercariae in each beaker were then counted under a dissecting microscope (10-40X magnification).

### Statistical Analysis

All statistical analyses were conducted with R (version 4.2.1). For one tank, the snail population crashed early in the experiment and was removed from all analyses. The first statistical analysis we conducted was a generalized additive model on adult snail abundance [mgcv package (Wood & Wood 2015), gam function, negative binomial error distribution, tank as a random intercept] to assess whether there were any nonlinearities through time. We discovered that snail abundance in all treatments showed a hump-shaped pattern through time (Fig. 1A), consistent with patterns in the field where snail abundances increase and decrease seasonally (Bakhoum *et al*. 2021b). Given that all treatments started with similar snail abundances and converged on similar abundances, we focused on weeks 7 through 15 for subsequent analyses because these were the majority of weeks where abundances seemed to diverge among the treatments. For this subset of data, we conducted generalized linear mixed effects models [package glmmTMB (Brooks *et al*. 2017)] that included the main effects of atrazine and fertilizer, an atrazine-by-fertilizer interaction, and tank as a random intercept. For response variables that were counts (adult and juvenile snails, egg masses, number of snails shedding, and cercariae shed per snail) and estimates of biomass (*C. demersum* mass, periphyton and phytoplankton abundance), we used negative binomial and Gaussian error distributions, respectively. Probability values were calculated using the Anova function in the car package (Fox *et al*. 2012) and figures were generated using ggplot2 (Wickham 2011). When there were significant main effects or interactions, we used the pairs function in the emmean package (Lenth 2023) to conduct a Tukey’s test to determine which pairs of the four treatments were significantly different. Finally, to test for correlations between plant biomass, phytoplankton, and periphyton, we standardized values (i.e., mean = 0, standard deviation =1 for each of the four treatments) and tested for correlations within rather than across treatments.

**Figure 1.**
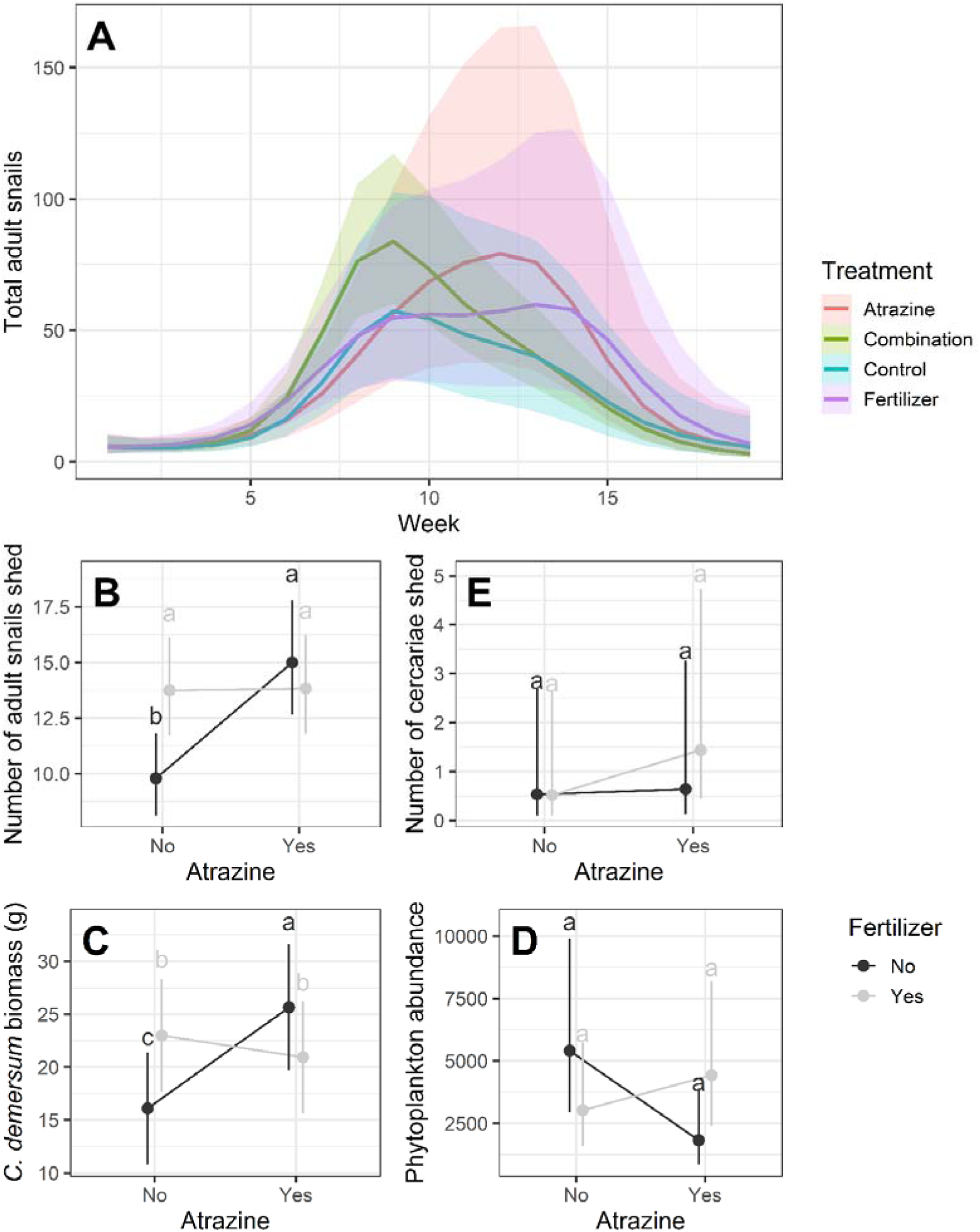
Effects of atrazine and fertilizer treatments on (**A**) adult snail abundance across 19 weeks of the experiment and on (**B**) adult snail abundance, (**C**) vegetation biomass, (**D**) phytoplankton abundance (measured as Ft, which represents raw fluorescence units or the intensity of the fluorescence signal detected by the instrument’s photodiode), and (**E**) *Schistosoma mansoni* cercariae shed per tank, averaged from weeks 7 to 15. In panel A, the lines are predictions from generalized additive mixed-effects model along with 95% confidence bands. This model indicated strong temporal dynamics with most deviations occurring between weeks 7 to 15. In panels B-E, points are predicted values (*n =* 5) with 95% confidence intervals and treatments that do not share a letter are significantly different based on a Tukey’s test.

## RESULTS

A significant interaction between atrazine and fertilizer was detected for our estimate of the number of adult snails (Table 1, Fig. 1B) and *C. demersum* biomass (Table 1, Fig. 1C). The control treatment had fewer adult snails and less *C. demersum* than atrazine alone, fertilizer alone, and the combination of atrazine and fertilizer, and these latter three treatments did not differ from one another for either response variable (Fig. 1B, C). The treatments had no detectable effects on periphyton or juvenile snails (Table 1, Fig. S1), but atrazine tended to reduce phytoplankton in the absence of fertilizer (Table 1, Fig. 1D) and fertilizer tended to increase egg mass production (Table 1). Although we had more adult snails in the tanks with atrazine or fertilizer relative to controls, we did not detect significantly more shedding snails or more cercariae shed from snails in these treatments. However, there were trends towards more of each in the atrazine plus fertilizer treatment (Table 1, Fig. 1E). Importantly, this might be because so few snails were infected and/or shed cercariae. Despite repeated miracidia exposure, most treatments averaged less than one snail and one cercariae shed per tank per week (Fig 1E). As predicted from the hypothesis that phytoplankton might shade submerged aquatic plants, phytoplankton abundance was negatively correlated with vegetation biomass (Fig 2).

**Figure 2.**
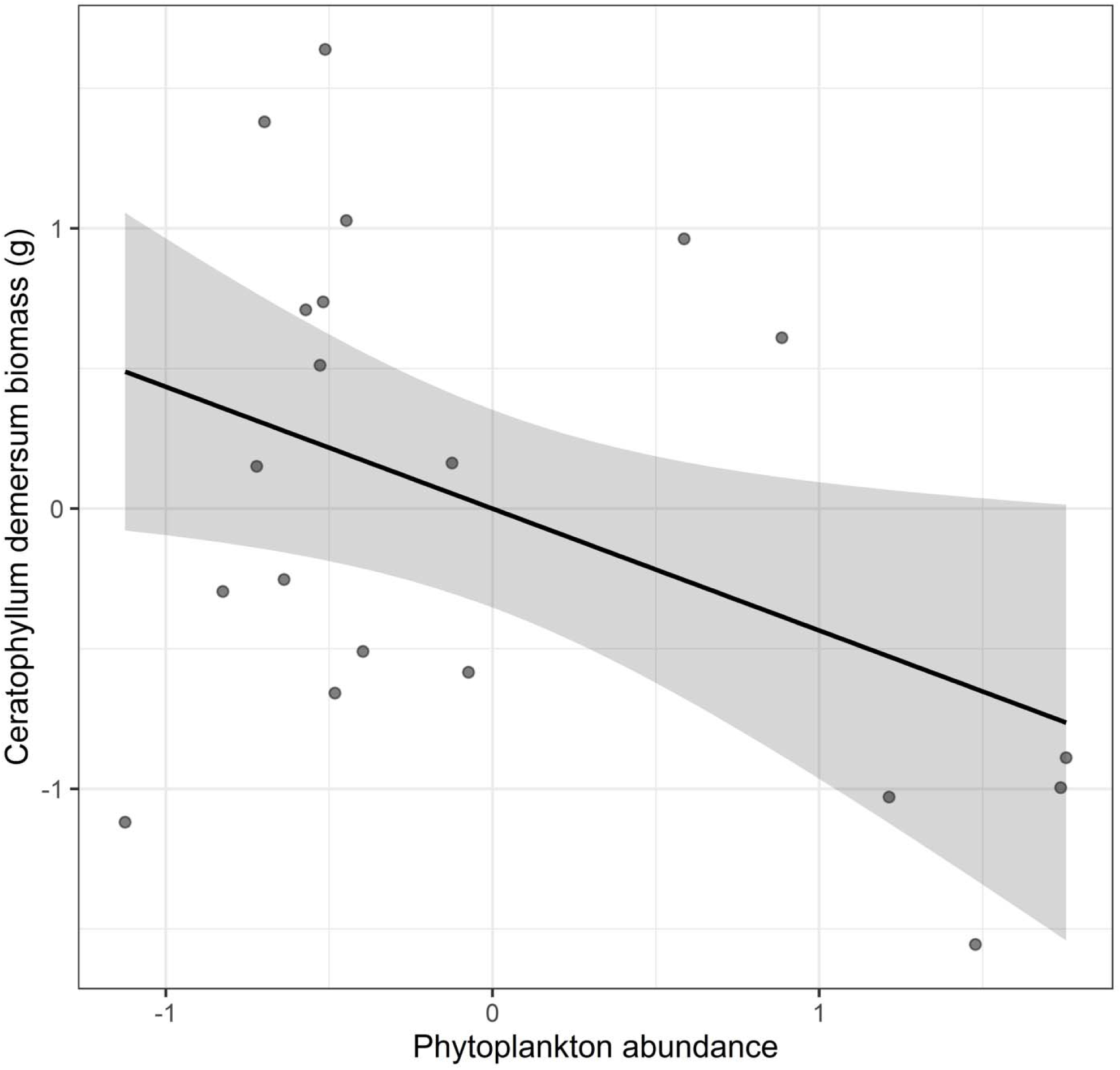
*Ceratophyllum demersum* biomass (natural log-transformed) is negatively correlated with phytoplankton abundance (measured as Ft, which represents raw fluorescence units or the intensity of the fluorescence signal detected by the instrument’s photodiode; *X^2^* = 4.66, *p* = 0.031). Shown are Z-score standardized data points within treatments, the best fit line, and a 95% confidence band.

**Table 1.** Results of ANOVA evaluating responses to atrazine, fertilizer, and their interaction. S.E.= standard error. Significant values are bolded.

|  | Atrazine |  |  | Fertilizer |  |  | Interaction |  |  |
| --- | --- | --- | --- | --- | --- | --- | --- | --- | --- |
|  | Coefficient<br>(s.e.) | Test<br>statistic<br>(X<sup>2</sup>) | *P* | Coefficient<br>(s.e.) | Test<br>statistic<br>(X<sup>2</sup>) | *P* | Coefficient<br>(s.e.) | Test<br>statistic<br>(X<sup>2</sup>) | *P* |
| Adult snails | 0.426 (0.127) | 5.211 | **0.022** | 0.339 (0.123) | 1.974 | 0.160 | -0.420 (0.171) | 6.054 | **0.014** |
| Juvenile snails | 0.056 (0.240) | 0.012 | 0.915 | -0.094 (0.229) | 0.594 | 0.441 | -0.074 (0.335) | 0.005 | 0.824 |
| Snail egg masses | -0.087 (0.763) | 0.904 | 0.341 | 0.778 (0.611) | 5.826 | **0.016** | 0.619 (0.888) | 0.486 | 0.486 |
| Infected snails | 0.225 (1.051) | 1.100 | 0.294 | 0.082 (1.042) | 0.660 | 0.417 | 0.819 (1.370) | 0.358 | 0.550 |
| Cercariae per tank | 0.184 (1.000) | 1.109 | 0.292 | -0.027 (1.000) | 0.525 | 0.469 | 0.836 (1.304) | 0.412 | 0.521 |
| Plant biomass | 9.552 (4.032) | 1.524 | 0.217 | 6.870 (3.801) | 0.262 | 0.609 | -11.593 (5.542) | 4.376 | **0.036** |
| Periphyton abundance | -0.015 (0.292) | 0.388 | 0.533 | -0.278 (0.296) | 0.397 | 0.529 | 0.287 (0.414) | 0.483 | 0.487 |
| Phytoplankton abundance | -1.085 (0.491) | 0.744 | 0.389 | -0.578 (0.444) | 0.055 | 0.814 | 1.464 (0.664) | 4.861 | **0.027** |

## DISCUSSION

Agricultural intensification is increasingly recognized as a major ecological driver of environmentally mediated infectious diseases, yet predicting how specific agricultural practices influence disease risk remains a central applied challenge (Rohr *et al*. 2019). Our results demonstrate that environmentally relevant applications of nitrogen–urea fertilizer and the herbicide atrazine, both singly and in combination, can restructure freshwater food webs in ways that favor key ecological drivers of schistosomiasis transmission. Specifically, agrochemical treatments increased snail intermediate host abundance and submerged aquatic vegetation while altering algal communities, consistent with bottom-up pathways that have been shown to regulate transmission risk across endemic regions (Wood *et al*. 2019; Haggerty *et al*. 2020; Haggerty *et al*. 2022; Rohr *et al*. 2023). Importantly, agrochemical mixtures produced non-additive effects on host and habitat dynamics, highlighting that the ecological consequences of agricultural pollution cannot be reliably inferred from single-compound exposures alone. Together, these findings underscore the value of applied ecological experiments for identifying mechanistic pathways through which realistic agrochemical regimes shape disease-relevant ecosystem processes.

Consistent with seasonal snail population dynamics in the wild where snails often start at low densities, peak, and then decline (Barbosa, Pimentel-Souza & Sampaio 1987; Bakhoum *et al*. 2021b), snail populations in our experiment remained at low levels for the first 6 weeks of the experiment before treatments exhibited considerable variation in abundance between weeks 7 to 15. Treatments ultimately re-converged as populations rapidly declined late in the experiment, perhaps because per capita resources declined once snails approached carry capacities that likely differed among treatments. Hence, we focused our analyses on the time periods where there were considerable abundances of snails in our tanks and separation among our treatments.

We originally hypothesized that fertilizer application alone would promote increases in snail biomass by increasing available nutrients for snail habitat and periphyton (Rohr *et al*. 2023). Our data showed that application of fertilizer alone did have observably positive effects on adult snail biomass, egg mass production, and vegetation biomass (Fig. 1). However, we did not detect a significant effect of fertilizer on periphyton (Fig. 1). Previous studies have shown that fertilizer strongly influences snail biomass by increasing periphyton, a food resource for snails (Halstead *et al*. 2018). However, this relationship can be challenging to observe because any increases in periphyton may have been quickly consumed by snails and converted to snail biomass (Civitello *et al*. 2018). Given that periphyton was the primary food resource, the fertilizer-driven increase in snail biomass was likely mediated by greater periphyton availability that snails rapidly consumed.

As predicted, atrazine alone had a positive effect on vegetation biomass (Fig. 1). This result is consistent with the hypothesis that atrazine disproportionately suppresses phytoplankton, suspended photosynthetic algae adapted to higher light conditions, relative to periphyton and submerged vegetation, which often thrive in lower-light environments (Guasch *et al*. 1998). By reducing phytoplankton, atrazine can indirectly promote the growth of aquatic vegetation and periphyton by increasing light penetration (Haggerty *et al*. 2022). Previous studies have shown that atrazine application reduces phytoplankton, leading to greater water clarity, nutrient availability, and periphyton productivity (Rohr *et al*. 2008; Staley, Rohr & Harwood 2011; Rohr, Halstead & Raffel 2012; Staley, Harwood & Rohr 2015; Rumschlag *et al*. 2019; Rumschlag *et al*. 2020). Our findings provide further support for this pathway because phytoplankton abundance was highest in controls and lowest under atrazine treatment, and phytoplankton was negatively correlated with vegetation biomass. Although we did not detect a significant relationship between phytoplankton and periphyton, atrazine increased adult snail abundance, consistent with the idea that snails quickly consumed any gains in periphyton, thereby potentially masking associations in our measurements. Overall, these results suggest that atrazine indirectly promotes snail populations by reducing phytoplankton shading and enhancing the growth of submerged vegetation and periphyton, key habitat and resources for snails.

Even though atrazine and fertilizer alone increased snail abundance and *C. demersum* biomass relative to controls, the combination of atrazine and fertilizer did not further increase snail abundance or *C. demersum* biomass relative to either treatment alone (Fig. 1). Plant biomass over weeks 7 to 15 was elevated relative to controls. While atrazine alone led to significantly more plant biomass than fertilizer alone, the effects of the herbicide-fertilizer mixture were not significantly different from either group. This supports the hypothesis that atrazine and fertilizer have a non-additive antagonistic interaction because fertilizer lessened the effects or facilitated the recovery of phytoplankton exposed to atrazine. This hypothesized mechanism for this antagonistic interaction was supported by the fact that phytoplankton abundance was higher in tanks with fertilizer and atrazine than atrazine alone.

Although parasite infection levels in snails were low in this experiment, this does not diminish the applied relevance of our findings. In schistosomiasis systems, transmission risk is more strongly constrained by the abundance of snail intermediate hosts and the availability of suitable habitat than by per-snail infection probability, particularly in early or low-intensity transmission settings (Civitello *et al*. 2018; Wood *et al*. 2019; Haggerty *et al*. 2020; Civitello *et al*. 2022). Ecological drivers such as submerged aquatic vegetation and host density regulate contact rates, parasite production, and spatial overlap with humans, and thus often precede detectable increases in parasite prevalence. Consequently, changes in host and habitat dynamics are widely used as leading indicators of transmission potential in both experimental and field contexts (Haggerty *et al*. 2020; Rohr *et al*. 2023). The consistent increases in snail abundance and vegetation observed here therefore represent biologically meaningful shifts in disease-relevant ecosystem processes, even in the absence of strong infection responses.

Previous studies have identified atrazine and fertilizer as primary drivers of elevated trematode infections in amphibians, with consistent effects across host–parasite taxa (Rohr *et al*. 2008; Schotthoefer *et al*. 2011; Rohr *et al*. 2015). In our experiment, we observed trends toward increased cercarial production in tanks receiving the atrazine–fertilizer mixture. Although these trends were non-significant, likely due to low infection levels across treatments, they are consistent with the broader literature showing that agrochemical-driven changes in host density and habitat can precede measurable increases in parasite output. Together with recent field evidence linking fertilizer use to aquatic vegetation, snail abundance, and human schistosomiasis risk (Rohr *et al*. 2023), these results reinforce the generality of agrochemical effects across trematode systems.

From a management and policy perspective, our findings have direct implications for agrochemical regulation in disease-endemic freshwater landscapes. Current risk assessment frameworks typically evaluate fertilizers and pesticides individually, yet our results show that common agrochemical mixtures can generate non-additive ecological effects on snail hosts and habitat that are central to schistosomiasis transmission (Halstead *et al*. 2014; Rohr, Salice & Nisbet 2016). Because snail abundance and submerged vegetation are tractable intervention points that respond predictably to environmental management, overlooking mixture effects may lead to systematic underestimation of disease risk associated with agricultural intensification (Rohr, Salice & Nisbet 2016; Rohr, Salice & Nisbet 2017). Incorporating mixture-based ecological responses into agrochemical risk assessment, alongside traditional toxicity endpoints, would better align regulatory decision-making with the ecosystem processes that ultimately govern transmission. Such an approach could help identify fertilizer–herbicide combinations that support crop production while minimizing unintended consequences for freshwater ecosystems and human health, advancing integrated strategies for sustainable agriculture and disease control in endemic regions (Halstead *et al*. 2014; Rohr, Salice & Nisbet 2017; Rohr *et al*. 2019; Haggerty *et al*. 2022).

Taken together, our findings show that common fertilizer–herbicide mixtures can restructure freshwater ecosystems in ways that favor ecological drivers of schistosomiasis risk. By increasing snail intermediate host abundance and submerged aquatic vegetation through bottom-up pathways, agrochemical inputs influence transmission-relevant processes that precede and constrain epidemiological outcomes (Wood *et al*. 2019; Civitello *et al*. 2022; Rohr *et al*. 2023). More broadly, this study illustrates how applied ecological experiments in semi-natural systems can reveal emergent ecosystem responses to realistic chemical mixtures that are not apparent from single-compound studies. Such mechanistic insight is essential for anticipating how agricultural intensification influences disease risk in freshwater ecosystems. Such mechanistic insight is essential for anticipating how agricultural intensification reshapes disease risk in freshwater ecosystems and for guiding applied ecological research that informs sustainable management in disease-endemic regions.

## Acknowledgements

This research was supported by funds from the National Science Foundation (DEB-2017785, DEB-2109293, BCS-2307944, and ITE-2333795), Frontiers Planet Prize, and the University of Notre Dame Poverty Initiative. The following reagent was provided by the NIAID Schistosomiasis Resource Center for distribution through BEI Resources, NIAID, NIH: *Biomphalaria glabrata*, Strain NMRI (Unexposed to *Schistosoma mansoni*), NR-21970.

## Author contributions

All authors designed the experiment and edited the manuscript. MRB, ES, MF conducted the research. JRR conducted the statistical analyses and developed the figures and tables. MRB and JRR wrote the initial draft. JRR acquired the funds for the study.

### Data availability statement

The data and code will be made available in Figshare upon acceptance.

### Conflict of interest statement

None of the authors have any conflicts of interests to declare.

---

**Supplemental Figure 1.**
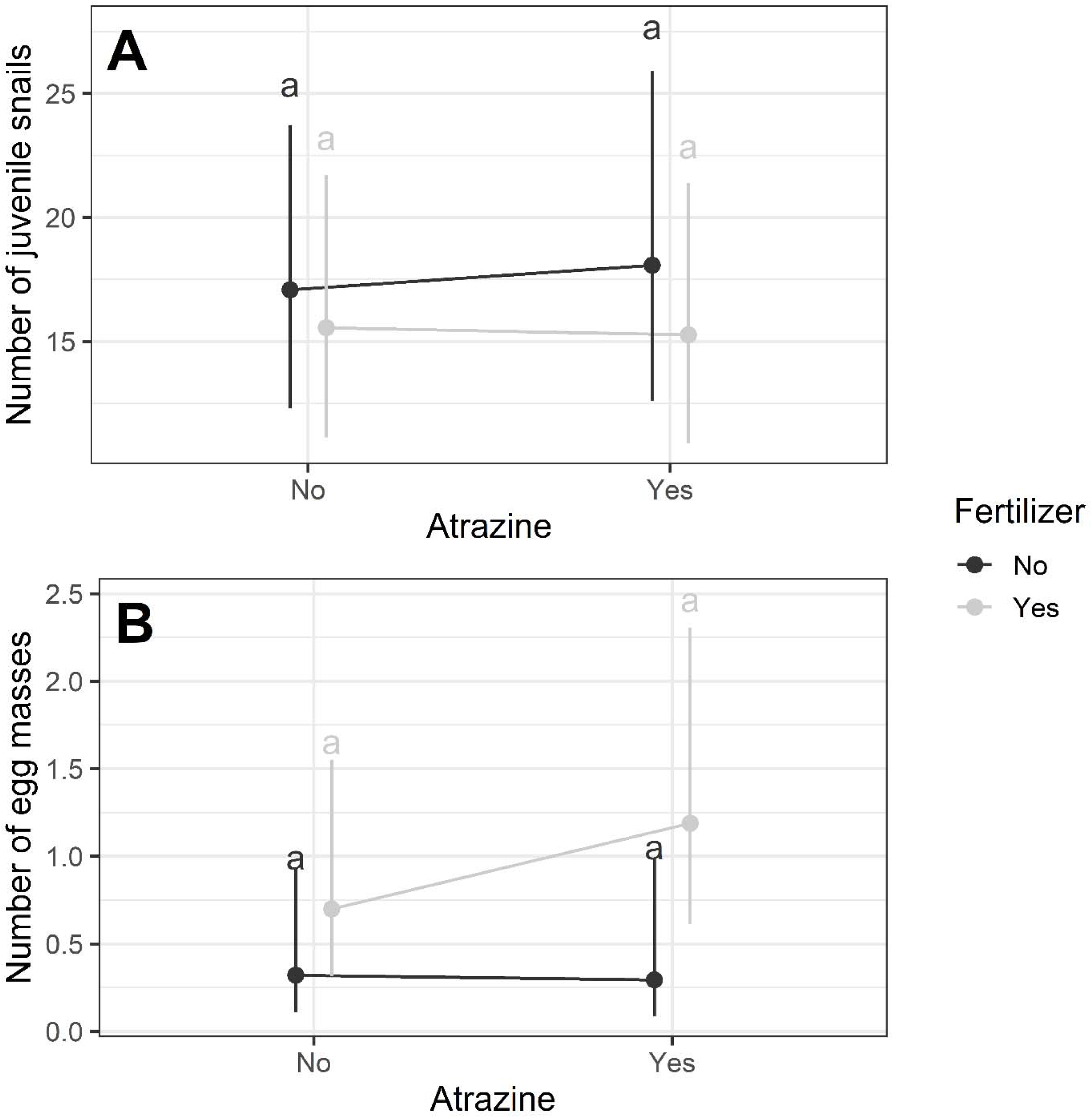
Effects of atrazine and fertilizer treatments on the average number of (**A**) juvenile snails and (**B**) egg masses per tank from weeks 7 to 15. Points are predicted values (*n =* 5) with 95% confidence intervals and treatments that do not share a letter are significantly different based on a Tukey’s test.

## Notes

### Competing Interest Statement

The authors have declared no competing interest.

